# Deploying genomics workflows on high performance computing (HPC) platforms: storage, memory, and compute considerations

**DOI:** 10.1101/2022.04.05.485833

**Authors:** Marissa E. Powers, Keith Mannthey, Priyanka Sebastian, Snehal Adsule, Elizabeth Kiernan, Jonathan T. Smith, Jessica Way, Beri Shifaw, David Roazen, Paolo Narvaez

## Abstract

Next Generation Sequencing (NGS) workloads largely consist of pipelines of tasks with heterogeneous compute, memory, and storage requirements. Identifying the optimal system configuration has historically required expertise in both system architecture and bioinformatics. This paper outlines infrastructure recommendations for one commonly used genomics workload based on extensive benchmarking and profiling, along with recommendations on how to tune genomics workflows for high performance computing (HPC) infrastructure. The demonstrated methodology and learnings can be extended for other genomics workloads and for other infrastructures such as the cloud.

## Introduction

Since the advent of Next Generation Sequencing (NGS), the cost of sequencing genomic data has drastically decreased, and the amount of genomic samples processed continues to increase [1,2]. With this growth comes the need to more efficiently process NGS datasets.

Prior work has focused on custom methods for deploying exomes on HPC systems [3], as well as best practices for deploying genomics workflows on the cloud [4]. This paper outlines how to optimize system utilization for one commonly used genomics workflow, along with recommendations on how to tune genomics workflows for HPC infrastructure.

The Broad Institute’s Genome Analysis Toolkit (GATK) Best Practices Pipeline for Germline Short Variant Discovery is a commonly used workflow for processing human whole genome sequences (WGS) datasets. This pipeline consists of 24 tasks, each with specific compute, memory, and disk requirements.

See S1 Table for a full list of the tasks and their requirements. Of these 24 tasks, six are multithreaded, and the rest are single threaded.

For multithreaded tasks, the genome is broken into shards, and each is executed as a parallel process. At the end of the task, the output datasets from all shards are aggregated and passed as a single input to the next task. This ability to “scatter” a task across multiple jobs, and then “gather” outputs for the next task is called “scatter-gather” functionality [5]. The ability to process multiple jobs concurrently is referred to as “parallelization.” Both scatter-gather functionality and parallelization are key concepts for efficiently distributing genomic pipelines on a system.

For example, in the task BWA, which aligns fragments output by the sequencer into a single aligned string, the genome is broken into 24 shards. On local high performance computing (HPC) infrastructure, each of these 24 shards is packaged as a single batch scheduler job. Once all 24 shards of BWA complete, the task MergeBamAlignment (Mba) consolidates the 24 output files into a single input file for the next task, MarkDuplicates, which is single threaded.

Each of these 24 BWA jobs is deployed on the cluster and executed in parallel, and each job is allocated a recommended four CPU threads and 14GB DRAM (see S1 Table). When BWA is running with these parameters, therefore, it consumes in total 96 threads and 336GB DRAM.

It’s important to note that while BWA can readily consume 96 threads on a system, most of the tasks are single threaded. Figure 1 below shows CPU utilization (gray) and memory utilization (in red) for the duration of the pipeline when processing a single 30X coverage human whole genome sequence (WGS). Note that memory utilization is close to its maximum for roughly only a third of the overall processing time. CPU utilization is at 100% for even less time.

**Fig 1.**
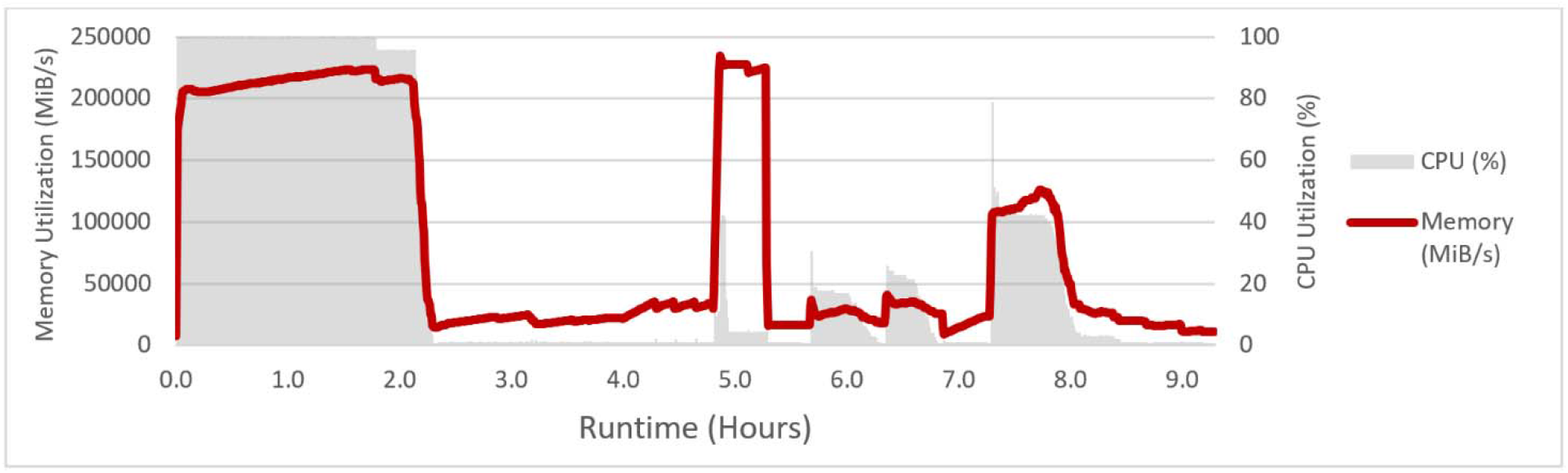
CPU and Memory Utilization for single WGS run. The runtime for key tasks in the pipeline (with start and end times in parens) is BWA (0.0-2.4); MarkDuplicates (2.4-4.8); SortSampleBam (4.8-5.7); BaseRecalibrator (5.7-6.3); GatherBQSRReports (6.3-6.4); ApplyBQSR (6.4-6.9); GatherBamFiles (6.9-7.3); HaplotypeCaller (7.3-8.7); MergeVCFs (8.7-9.3). CPU Utilization is at 100% for BWA, and 40-50% for HaplotypeCaller. Memory utilization is close to 100% only for BWA and SortSampleBam. Because of this heterogeneity in resource utilization, achieving maximum throughput requires efficient scheduling of multiple WGS samples in parallel.

Because of this heterogeneity, making efficient use of HPC infrastructure requires tuning and orchestration of the workflow. First, these tasks need to be efficiently sharded and distributed across the cluster. Second, each task needs to be allocated the optimal number of threads and memory. Local disk must be used for temporary storage. Finally, the tasks need to take advantage of underlying hardware features.

This paper outlines the impact of each of these factors on performance, and details best known methods for configuring the Germline Variant Discovery pipeline on local HPC infrastructure.

## Materials and Methods

Benchmarking was performed on a five-server cluster, with one application server and four compute servers. A full hardware configuration can be found in Supporting Information S2 Table.

The publicly available NA12878 30X coverage whole genome sequence (WGS) dataset was used for all benchmarking. For the resource profiling (e.g. Fig 1), a single sample was run. For throughput tuning 40 WGS were submitted concurrently.

Jobs were orchestrated on the cluster with Slurm. The Broad Institute provides a Slurm backend for Cromwell, which can be found in the Cromwell documentation [6]. GATK Best Practices Pipelines are defined in Workflow Description Language (WDL). A given WDL defines which GATK tools to call in the form of tasks and is accompanied by a JSON file with dataset locations and other configuration settings.

The full list of tasks in the Germline Variant Discovery pipeline can be found in S1 Table, along with the compute requirements for each task. Testing was performed with GATK v4.1.8.0. A full software bill of materials is provided in the Supporting Information S3 Table.

The specific WDL and JSON files used for testing can be found in S3 Table. The recommended resource allocation values are included in those workflows.

## Results

### 1. Tuning Resource Allocation Values

To allocate specific amounts of cores and memory to each task, the HPC batch scheduler must be configured to enable consumable resources. With Slurm, this is set in slurm.conf by specifying both “SelectType=select/cons_res” and “SelectTypeParameters=CR_Core_Memory.” Additional detail on consumable resources is available in the Slurm documentation [7].

Another key component of resource tuning is Hyperthreading. With Hyperthreading turned on (HT=On) two processes can be executed simultaneously on a single physical core. Testing shows a 10% overall pipeline speedup with HT=On. With HT=On, a task is allocated a set number of threads, with two threads available per physical core.

The most compute-intensive task in the Germline Variant Discovery pipeline is BWA. Manipulating the number of threads per shard for BWA has a substantial impact on the overall runtime of the pipeline, as shown in Figure 2A.

**Fig. 2.**
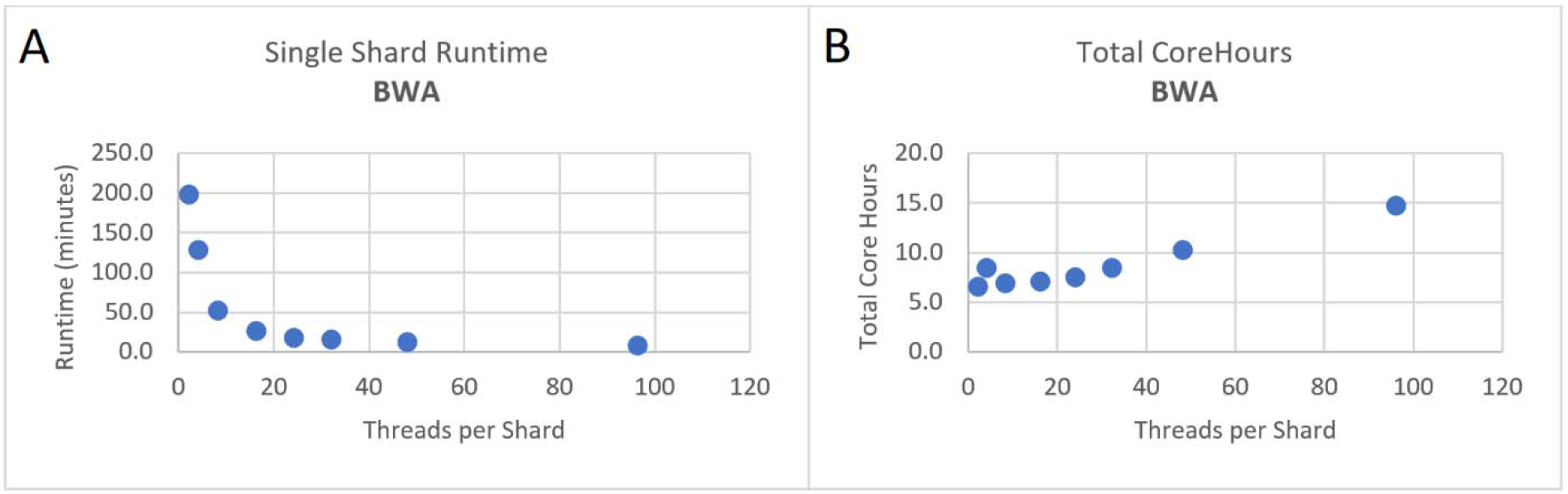
Impact of Increasing threads-per-shard on BWA Performance. Figure 2A shows the effect of increasing threads per shard on single shard runtime for BWAmem. Figure 2B shows the effect of increasing the number of threads per shard on total corehours consumed.

Figure 2A shows impact of increasing the number of threads per shard on the shard runtime. When two threads are allocated per shard, the runtime is 200 minutes. Increasing to 16 threads per shard decreases the runtime by 10X down to 27 minutes.

Increasing the thread count higher than 16 threads per shard has limited positive impact on shard runtime. Figure 2B shows the impact, however, on total corehours. Increasing the number of threads per shard gradually increases the total corehours consumed on the cluster. When processing a single WGS, 16 threads per shard provides a fast task runtime while limiting corehours.

S1 Table provides a detailed list of thread count and memory recommendations for each individual task in the pipeline, including BWA. These values were determined empirically specifically optimizing for throughput processing. The Discussion covers considerations when optimizing for fastest single sample runtime, as well as considerations for cloud infrastructure.

While 16 threads per shard results in the fastest single shard runtime, and the fastest runtime for BWA, it does not necessarily result in the best throughput, or number of genomes that can be processed on a system per day. On a 4-server 2-socket system with 24-core CPUs and HT=On, there are 384 available threads available at any given time. Setting BWA to consume 16 threads per shard for 24 shards results in BWA consuming all 384 threads for a single WGS. Adjusting this thread count to, for example, 4 threads per shard, results in a longer runtime for BWA but allows for processing four WGS samples in parallel.

While this section has focused on BWA tuning, similar methods were used to identify optimal thread and memory allocations for each task in the pipeline. These recommended values can be found in S1 Table.

### 2. Distributing Tasks Efficiently

The second most compute-intensive task in the pipeline is HaplotypeCaller, which performs variant calling. For HaplotypeCaller, the number of shards the tasks are distributed across is set in the WDL as the variable “scattercount.”

Figure 3 shows the relationship between the runtime of HaplotypeCaller and scattercount. When HaplotypeCaller is sharded into just two jobs (scattercount=2), the task takes 400 minutes to complete. As shown in Figure 3A, the task runtime decreases as scattercount increases up to scattercount=48. Beyond scattercount=48, there’s limited benefit in further sharding the task into smaller jobs.

**Figure 3.**
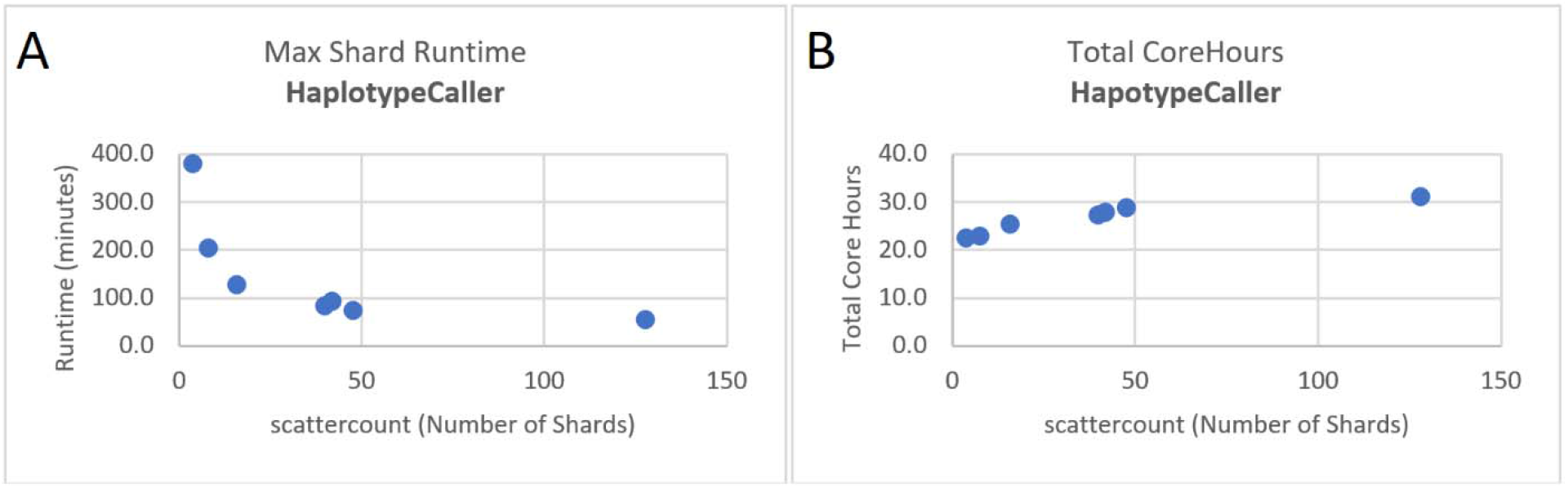
Impact of scattercount on HaplotypeCaller Runtime and total Core Hours. As scattercount increases, the runtime of the longest running shard decreases (A) while the total corehours consumed increases (B).

Figure 3B shows the relationship between scattercount and total corehours consumed by HaplotypeCaller. Corehours gradually increases with scattercount. As HaplotypeCaller is split into more small jobs the total corehours consumed increases.

It’s important to note that scattercount cannot be arbitrarily set without considering potential artifact generation. For this reason, scattercount is set specifically to 48. Concordance analysis is always required when tuning scattercount to ensure fidelity.

### 3. Local Disk for Temporary Storage

While BWA and HaplotypeCaller are the two most compute-intensive tasks in the pipeline, one of the longest running tasks is the single threaded MarkDuplicates. MarkDuplicates takes in a BAM or SAM file and compares and identifies duplicate reads.

The uncompressed files processed by MarkDuplicates for a 30X human whole genome sequence can total over 200GB in size. The task is highly dependent on fast local storage for processing these datasets.

Figure 4 shows the impact of running with a local Solid State Drive (SSD) compared to running without an SSD and just using the parallel file system. With an SSD, MarkDuplicates runtime is 2.5 hours. Without an SSD, MarkDuplicates runtime is 37.6 hours.

**Fig. 4.**
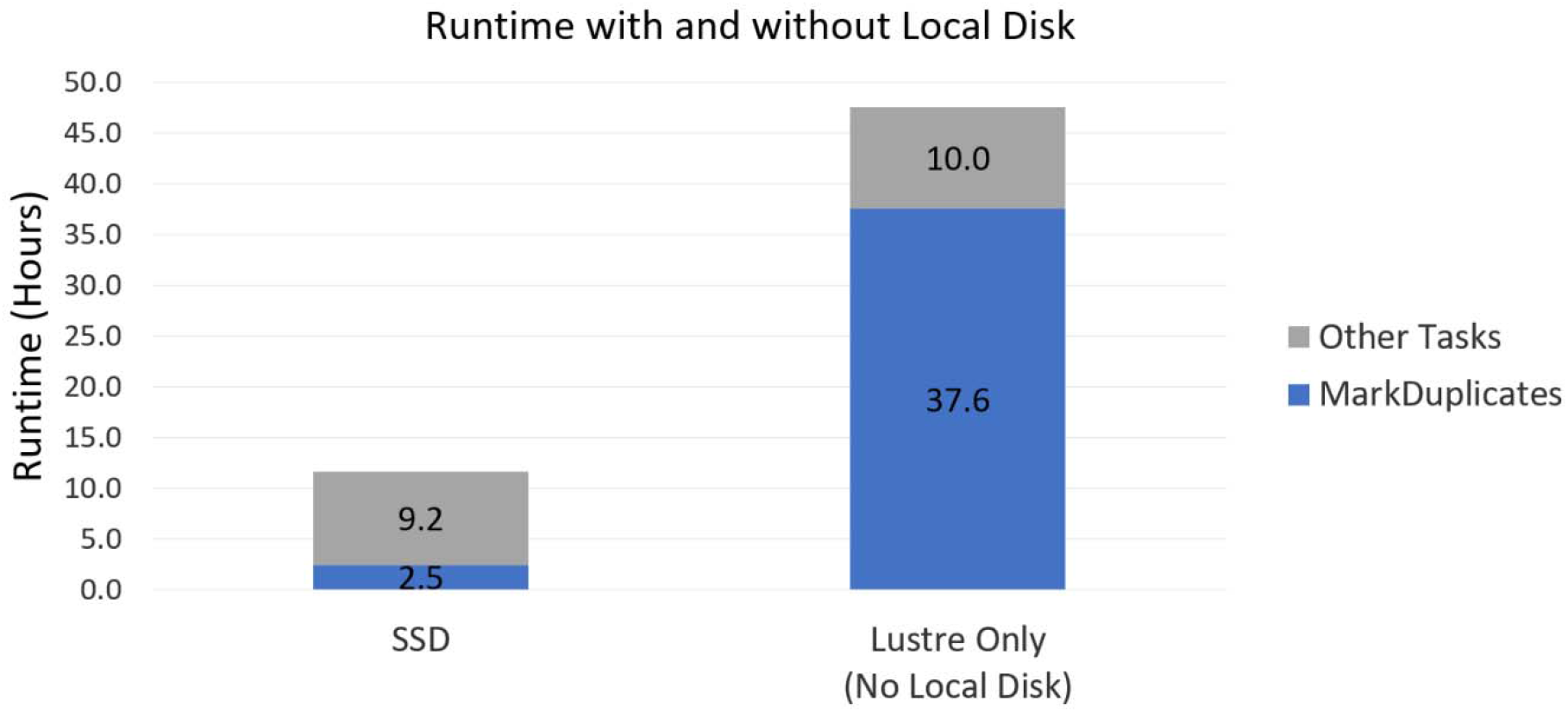
Local disk for MarkDuplicates Performance. The runtime for the entire Germline Variant Calling pipeline drastically decreases with use of a local SSD. This is primarily due to the decrease in runtime of MarkDuplicate (blue).

S2 Table includes an NVMe Pxxx SSD in each of the compute servers, largely for the sake of MarkDuplicates processing.

### 4. Genomics Kernel Library (GKL) to Utilize Hardware Features

Ultimately each shard of each task is executed on a single thread of a CPU. Ensuring these tasks are able to take advantage of the underlying CPU features is a key factor for performance.

As of GATK 4.0, a number of tasks in the Germline Variant Discovery pipeline have been accelerated to take advantage of Intel AVX-512 Instructions through the Genomics Kernel Library (GKL). GKL is developed and maintained by Intel and is distributed open source with GATK [8].

GKL includes compression and decompression from Intel’s ISA-L and zlib libraries, as well as AVX-512 implementations of PairHMM and Smith-Waterman [8-10]. PairHMM and Smith-Waterman are two key kernels included in a number of genomics tasks, including HaplotypeCaller.

Figure 5 shows the benefit of GKL compression for the three tasks with the largest input file sizes: ApplyBQSR (133GB), GatherBamFiles (62GB) and MarkDuplicates (222GB). GKL provides compression at levels from 1-9 (CL=1-9). CL=1 (orange) with GKL provides a 2-4X compression ratio relative to with no compression (blue). The compression ratio continues to improve as compression levels increase up to level 5.

**Fig. 5.**
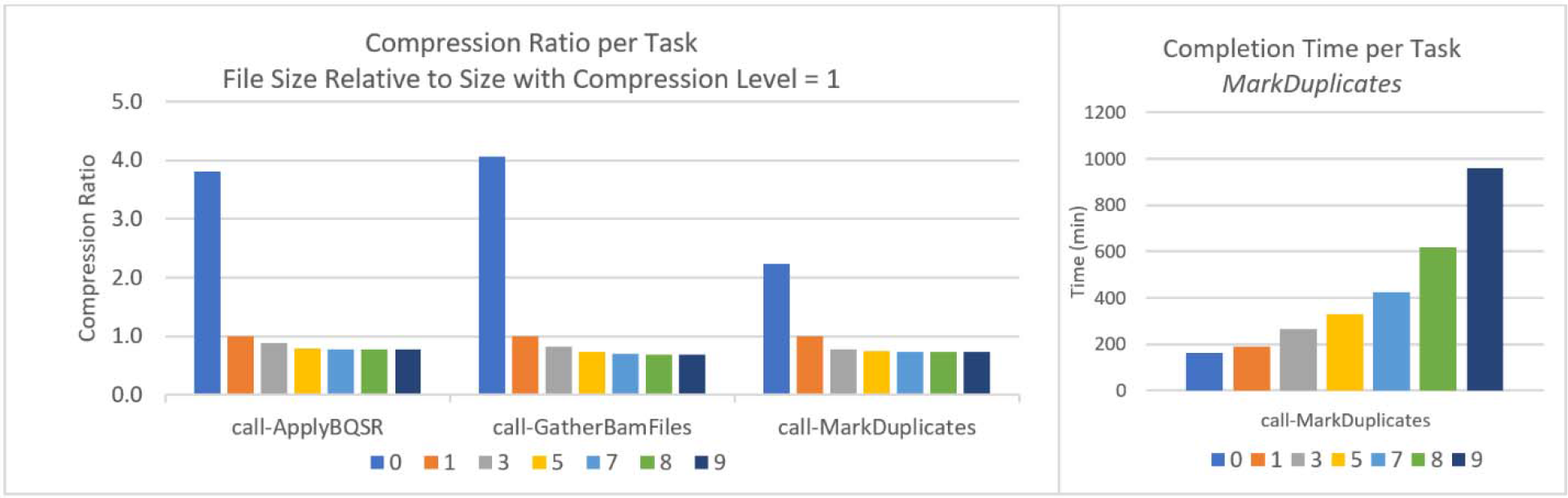
Compression with GKL for tasks with largest input file sizes. The Genomics Kernel Library (GKL) performs compression at levels 1 through 9 (CL=1-9). Compression ratios relative to CL=1 are shown in Figure 5A for three different tasks in the pipeline. Figure 5B shows the impact of each compression level on completion time for MarkDuplicates.

Figure 5B shows the task runtime as a function of compression level for these three tasks.

As the name suggests, MarkDuplicates checks the input BAM file for duplicate reads, and tags any identified duplicates [11]. In doing so the task reads and writes small (kB) intermediary files throughout the 2+ hours of processing. Each of these intermediary files is compressed and decompressed. Because of this, higher compression levels result in a high runtime cost with this task (see Figure 5B).

Based on these results, compression level is set to CL=2 in GATK 4.2.0.0. This compression level provides a good balance between high compression ratio across tasks (Figure 5A) and low runtime for MarkDuplicates (Figure 5B).

Figure 6 shows the difference between HaplotypeCaller runtime with the AVX-512 implementations of both kernels (left) compared to with the original implementations with no AVX instructions (right). The middle bar shows the runtime with the AVX512 implementation of SW and the Java AVX2 implementation of pairHMM.

**Fig. 6.**
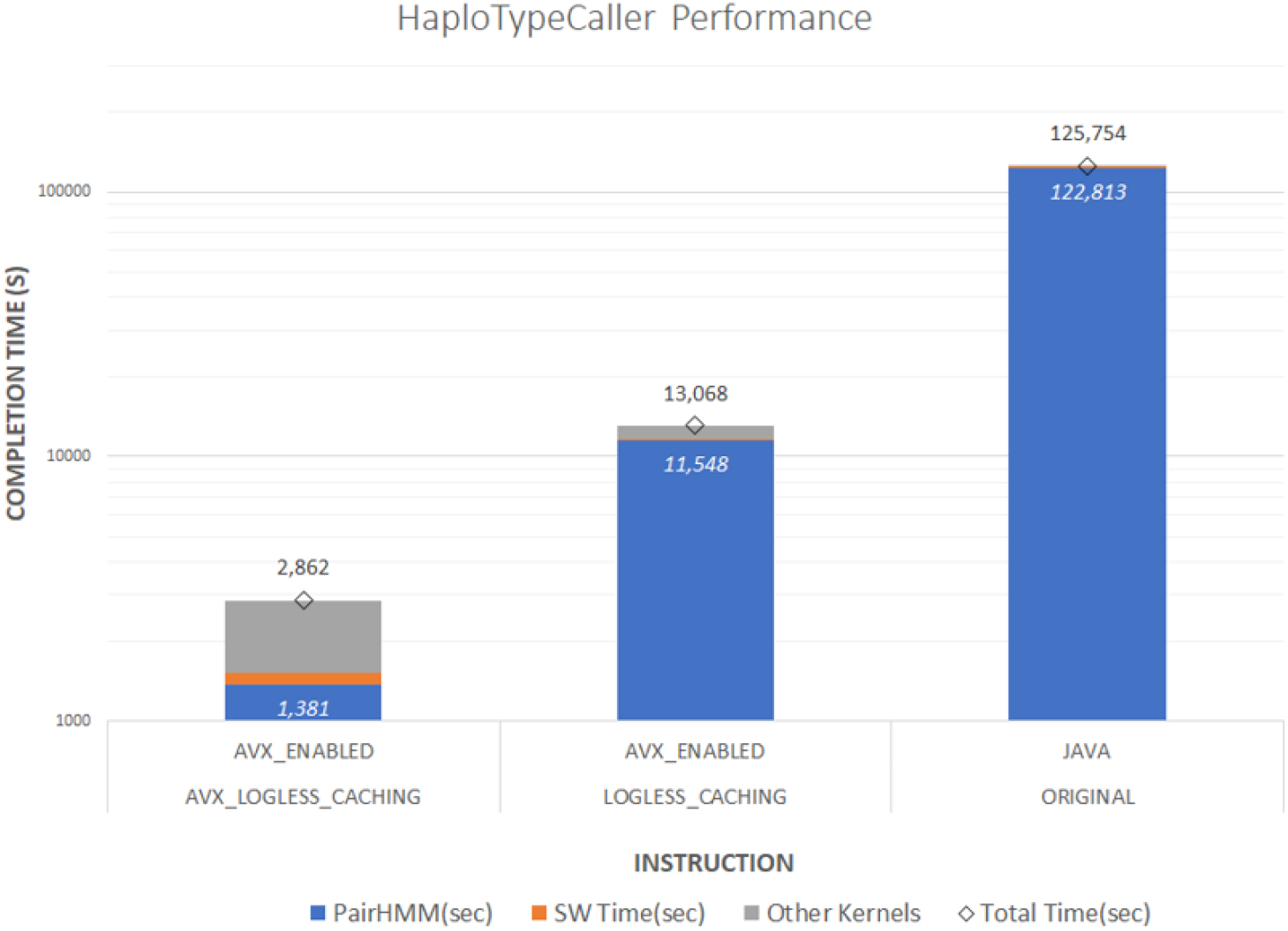
Impact of GKL AVX flags on HaplotypeCaller Performance. HaplotypeCaller performance, as measured by task runtime in seconds, drastically improves with the use of GKL pairHMM and SmithWaterman (SW). Runtimes for pairHMM are shown in blue; SW in orange.

Note the y-axis is a log scale. Without the AVX implementations, HaplotypeCaller takes 125,754 seconds, or 35 hours, to complete. With the GKL AVX512 implementations the same task completes in less than one hour.

Notably, users do not need to set any special flags to run with the GKL implementations. As shown in the HaplotypeCaller documentation, running with default flag (FASTEST_AVAILABLE) automatically detects if the underlying CPU includes support for AVX-512 instructions and, if so, deploys the GKL implementation [12].

## Discussion

As illustrated above, optimal performance of the Germline Variant Discovery pipeline is dependent on (1) efficiently distributing tasks across the cluster; (2) tuning resource allocation values; (3) utilization of fast local storage; and (4) libraries that take advantage of underlying CPU features.

Intel and Broad Institute have partnered to form the Intel-Broad Center for Genomics Data Engineering. The Genomics Kernel Library (GKL) is a direct outcome of this joint engineering Center. As part of this partnership, many of the configuration recommendations outlined above (eg the Slurm backend for Cromwell and resource allocation values) are directly incorporated into the Broad Institute workflows and documentation.

A specific reference architecture, including a detailed Installation Guide, is available as the Intel Select Solution for Genomics Analytics [13]. The Solution includes a detailed hardware and software configuration similar to that provided in the Supporting Information (A2 and A3).

The recommended resource allocation values provided in S1 Table have been specifically tuned for throughput, or the number of WGS samples that can be processed on a cluster per day. For institutions processing dozens, hundreds, or even thousands of samples per month, this throughput metric is a higher priority than single sample processing time. In these scenarios, reducing the number of threads per task allows for more jobs to run concurrently on the system, increasing throughput.

In other cases, such as with single threaded tasks, it makes sense to increase the thread allocation to increase throughput. As shown in S1 Table, most tasks in the pipeline are allocated two threads each despite being single threaded. This is specifically to optimize for throughput. The second thread (1) allows for Java collection and (2) intentionally limits the number of jobs concurrently running on the system. Limiting the number of overall jobs helps ensure each task has sufficient memory while also leaving sufficient memory for scheduling and system level operations.

For scenarios where single sample processing time is the highest priority, increasing the threads and memory allocated per task will reduce single sample runtime, while decreasing the overall throughput of the cluster. A workflow optimized for single sample runtime is provided in the same repository as the throughput WDL (see S3 Table).

Increasing threads per task is also beneficial in the cloud. When deploying the Germline Variant Discovery pipeline through the Broad Institute’s Platform as a Service (PaaS) Terra.bio, each shard of each task is allocated its own VM with a set number of virtual CPUs (vCPUs) and DRAM. In this scenario, each of the 24 BWA shards is allocated 16 vCPUs, compared to the four threads per shard recommended for local deployments. Allocating each 16 vCPUs to each BWA shard does not negatively impact the runtime of other tasks and samples on the cloud, since there are no infrastructure scale constraints. Van der Auwera and O’Connor provide a detailed guide on best practices for deploying Broad Institute workflows on the cloud [4].

As shown, the optimal workflow configuration is dependent on both underlying infrastructure and key performance metrics (e.g. throughput vs single sample runtime). Profiling workloads with methods described here can be extended to genomics workflows beyond Germline Variant Calling. Future work will include tuning additional workflows as well as comparing cloud and local performance considerations.

## Key Points

- Because genomics workflows consist of pipelines of tasks with heterogeneous compute requirements, achieving maximum throughput requires efficient tuning and orchestration of these workflows.
- Tasks need to be efficiently sharded and distributed across the cluster; Each task needs to be allocated the optimal number of threads and memory; and local disk must be used for temporary storage.
- The Genomics Kernel Library (GKL) improves GATK performance by taking advantage of AVX-512 instruction sets and accelerating compression and decompression.

## Acknowledgements

The authors thank Michael J. McManus PhD for his input and guidance from study conception through data analysis. We thank Kyle Vernest, Kylee Degatano, and Louis Bergelson for technical support throughout benchmarking.

## Supporting Information

**S1 Table. Recommended Resource Allocations for Germline Variant Discovery tasks.**

**S2 Table. Hardware Configuration Used for Testing.**

**S3 Table. Software Configuration Used for Testing.**

