## Supplementary Tables for "Deploying genomics workflows on high performance computing (HPC) platforms: storage, memory, and compute considerations"

**S1 Table.**  **Recommended Resource Allocations for Germline Variant Discovery tasks.**

| Task | *Shards* | Threads  *per Shard* | Physical Memory *per Shard* | JavaHeap Memory *per Shard* |
| --- | --- | --- | --- | --- |
| ApplyBQSR | *19* | 2 | 4GB | 3g |
| BaseRecalibrator | *18* | 2 | 6GB | 4g |
| CalculateReadGroupChecksum | *1* | 2 | 2GB | 1g |
| CheckContamination | *1* | 2 | 2GB |  |
| CheckFinalVcfExtension | *1* | 1 | 2GB |  |
| CheckFingerprint | *1* | 2 | 1GB | 1g |
| CollectAggregationMetrics | *1* | 2 | 7GB | 5g |
| CollectGvcfCallingMetrics | *1* | 2 | 3GB | 2g |
| CollectQualityYieldMetrics | *1* | 1 | 2GB | 0.128g |
| CollectRawWgsMetrics | *24* | 2 | 3GB | 2g |
| CollectReadgroupBamQualityMetrics | *24* | 2 | 7GB | 5g |
| CollectUnsortedReadgroupBamQualityMetrics | *1* | 2 | 7GB | 5g |
| CollectWgsMetrics | *1* | 2 | 3GB | 2g |
| ConvertToCram | *1* | - | 3GB |  |
| CramToBam | *1* | - | 3GB |  |
| CreateSequenceGroupingTSV | *1* | 2 | 2GB |  |
| CrossCheckFingerprints | *1* | 2 | 2GB | 2g |
| GatherBamFiles | *1* | 2 | 3GB | 2g |
| GatherBqsrReports | *1* | 2 | 3GB | 3g |
| GetBwaVersion | *1* | 1 | 1GB |  |
| HaplotypeCaller | *50* | 2 | 7GB | 6g |
| MarkDuplicates | *1* | 2 | 7GB | 4g |
| MergeVCFs | *1* | 2 | 3GB | 2g |
| SamToFastqAndBwaMemAndMba | *24* | 4 | 10GB | 2g |
| ScatterIntervalList | *1* | 2 | 2GB | 1g |
| SortSampleBam | *1* | 40 | 5GB | 200GB |
| ValidateBamFromCram | *1* | 2 | 7GB | 6g |
| ValidateGVCF | *1* | 2 | 4GB | 3g |

**S2 Table. Hardware Configuration Used for Testing.**

|  | Component |
| --- | --- |
| System | 4 compute nodes |
| CPU (per node) | 2x Intel® Xeon® Platinum 6252 Processor (24 cores, 2.1GHz) |
| Hyperthreading | On |
| Turbo | On |
| Memory (per node) | 384 GB (12 slots; 32GB per slot; 2933 MHz) |
| Boot Drive (per node) | 1x Intel 960GB SSD OS Drive |
| Local Disk (per node) | 1x P4610 1.6TB 3DNAND PCIe |
| Network Interface Card (NIC) | 1x Intel X722 |
| Filesystem | Lustre Parallel File System |

**S3 Table. Software Configuration Used for Testing.**

|  | Component |
| --- | --- |
| OS | CentOS 7.6 |
| Workload | GATK Best Practices Pipelines for Germline Variant Calling  [*https://github.com/gatk-workflows/intel-gatk4-germline-snps-indels*](https://github.com/gatk-workflows/intel-gatk4-germline-snps-indels) |
| Tools | GATK 4.1.8.0, BWA-0.7.17 (r1188), Picard 2.16.0-SNAPSHOT, samtools 1.9 |
